# Benchmarking microRNA Target Prediction Algorithms Using Single-Cell Co-Sequencing Data

**DOI:** 10.1101/2025.06.23.661076

**Authors:** Louise Velut, Laura Fancello, Nadia Cherradi, Laurent Guyon

## Abstract

**Background:** MicroRNAs (miRNAs) are small non-coding RNAs that play pivotal roles in the post-transcriptional regulation of gene expression, influencing a wide range of physiological and pathological processes. Accurately identifying miRNA targets is crucial for understanding miRNA modes of action. To this aim, a plethora of algorithms have been developed to predict miRNA targets, each employing distinct methodologies and relying on different features. The limited overlap among target predictions generated by various algorithms underscores the necessity for comprehensive and independent benchmarks to evaluate their performance.

**Methods:** We selected seven algorithms among the most popular ones to perform a benchmark with an original approach using recently published datasets of miRNA-mRNA co-sequencing at the single-cell level. We used Gene Set Enrichment Analysis to assess algorithm’s capabilities to predict sets of targets statistically anti-correlated in expression with miRNAs. We worked with both co-sequencing datasets of human and mouse single-cells.

**Results:** Our benchmark shows high performances for mirDIP, which corresponds to the consensus result of 24 different algorithms, for human miRNAs. In human cell lines, Diana microT, TargetScan, miRmap, and miRDB also provide excellent results, while RNA22 and miRWalk exhibited poorer results. In mouse primary cells, Diana microT leads, closely followed by miRmap. Intriguingly, RNA22 performs better in mouse primary cells than in human cell lines. Moreover, our benchmark highlights the benefit of reducing targets to the experimentally validated ones. Finally, we demonstrated that performance varies depending on the number of targets used, with TargetScan performing better than mirDIP when considering only a few dozen targets.

## 1. Introduction

MicroRNAs (miRNAs) are small non-coding RNAs that function as key post-transcriptional regulators of messenger RNA (mRNA) expression. Their regulatory activity is implicated in a wide spectrum of biological processes, encompassing both physiological homeostasis and pathological conditions^1^.

Accurately identifying miRNA targets is essential for understanding how miRNAs function and for utilizing them as biomarkers or including them in future therapeutics. Various experiments aimed at identifying miRNA targets may not yield sufficient results due to the different cellular conditions that can influence the effects of miRNAs on their targets. To enhance the identification of miRNA targets, many algorithms have been developed, each using distinct methodologies and relying on different features. The limited overlap among the target predictions generated by these algorithms, as well as with experimentally validated targets, highlights the complexity of this task. This emphasizes the need for comprehensive and independent benchmarks to evaluate the performance of these prediction methods.

In this study, we focused on establishing a benchmark of seven widely recognized miRNA target prediction algorithms: Diana microT^2^, miRDB^3^, mirDIP^4^, miRmap^5^, miRWalk^6^, TargetScan^7^, and RNA22^8^.

Additionally, we utilized two experimentally validated databases of miRNA targets, miRTarBase^9^ and TarBase^10^. The benchmark is based on an innovative methodology that utilizes recently published datasets of miRNA-mRNA co-sequencing obtained at the single-cell level. Specifically, we employed co-sequencing datasets derived from four human cell lines as well as single cells from mouse lung tissue. These datasets present unique opportunities to investigate the relationships between miRNAs and their targets and have not been previously employed for the development of any existing predictive algorithms. Our objective is to evaluate the performance of the selected algorithms and databases in identifying targets that exhibit a statistically significant inverse correlation with miRNA expression. To this aim, we used Gene Set Enrichment Analysis (GSEA)^11^ to investigate miRNA-targets correlations enrichment.

The aim of this work is to effectively benchmark prediction algorithms and experimentally validated databases of miRNA targets using independent datasets. This approach enables to provide clear recommendations on which tools to use, depending on the species in question and the expected number of targets. Additionally, we intend to share our code, allowing users to incorporate their preferred algorithms into our benchmark.

## 2. Materials and methods

### microRNA-mRNA co-sequencing datasets

We analyzed publicly available single-cell miRNA-mRNA co-sequencing datasets from human cell lines and mouse lung cells, originally published by Li *et al*.^12^. After filtering, we included cells that had more than 5,000 expressed mRNAs and less than 1.4% mitochondrial gene expression. This resulted in the use of 523 K562 cells, 467 293T cells, 590 A549 cells, and 477 HeLa cells. For the murine datasets, all 9,403 available cells were included in the analyses.

Normalization of miRNA and mRNA expression values was performed using Seurat with default parameters. For the benchmark analysis, we set a minimum mean expression threshold for each cell type to select the relevant miRNAs. This threshold was determined using the mean expression estimated through LOESS (Locally Estimated Scatterplot Smoothing) at the point where the standard deviation was maximized. All expressed mRNAs were included in the subsequent analyses.

### microRNA target prediction algorithms and experimentally validated target databases

We selected seven algorithms from the most popular ones, each of which has been cited more than 200 times since 2020, and for which predictions were easily downloadable from their respective websites. Diana-microT^2^ from 2023 predictions were downloaded from the website: https://dianalab.e-ce.uth.gr/microt_webserver/#/. miRDB^3^ v6.0 predictions were downloaded from the website: https://mirdb.org/. mirDIP^4^ v5.2 predictions were downloaded from the website: https://ophid.utoronto.ca/mirDIP. miRmap^5^ 1.2 predictions were downloaded from the website: https://mirmap.ezlab.org/. miRWalk^6^ v3 predictions were downloaded from the website: http://mirwalk.umm.uni-heidelberg.de. RNA22^8^ v2 predictions were downloaded from the website: https://cm.jefferson.edu/rna22-full-sets-of-predictions/. TargetScan^7^ v8.0 predictions were downloaded from the website: https://www.targetscan.org/vert_80/.

Experimentally validated target lists from miRTarBase^9^ 2025 and TarBase^10^ v9 were respectively downloaded from http://mirtarbase.cuhk.edu.cn/~miRTarBase/miRTarBase_2025 and https://dianalab.e-ce.uth.gr/tarbasev9.

### Targets selection

For our analyses, we selected “top targets” based on the interaction scores predicted by algorithms for each miRNA-target pair. For a given “top n,” we used the targets with the best scores within the range of n ± 10%. If a target appeared multiple times in an algorithm’s predictions with different scores, only the best score was considered for selection.

In cases where scores were tied, we either included all targets sharing the same score or excluded all of them. If it was not possible to select n ± 10% of the targets due to a lack of predicted targets or ties in scores, we did not select any targets for that particular miRNA and, consequently, did not conduct the corresponding analysis.

It is important to note that experimentally validated target databases do not provide assigned interaction scores for each pair. To facilitate “top target” analyses, we used scores from mirDIP for human miRNAs and from Diana microT for murine miRNAs. Only targets with assigned scores were included; experimentally validated targets that were not predicted by mirDIP or Diana microT were excluded from these “top target” analyses.

### Correlation analyses of miRNA-mRNA pairs

We utilized Gene Set Enrichment Analysis (GSEA) ^11^ to evaluate the algorithm’s ability to predict sets of targets that exhibit statistically anti-correlated expression with miRNAs. For each miRNA of interest, we ranked all mRNAs based on the Pearson correlation between their expression levels and those of the miRNA. For each list of targets, we calculated an Enrichment Score (ES), which is negative if the correlations with targets are significantly enriched toward negative values and positive if enriched toward positive values. Each enrichment is accompanied by a p-value, which is presented on a log10 scale in the figures. We conducted GSEA using the clusterProfiler package in R^13^.

### Performance evaluation

A performance score was assigned to each algorithm to evaluate its ability to predict targets that are anti-correlated with miRNAs. For each target number condition and cell type, the performance score for an algorithm was calculated as the ratio of miRNAs that showed significant negative enrichment to the maximum number of miRNAs with significant negative enrichment identified by any algorithm. This calculation produced a score ranging from 0 to 1 for each algorithm and cell type. These scores were then summed, resulting in a total score of up to 4 for human cell lines and up to 22 for mouse lung cell types. The overall performance score for each algorithm was determined as the average of its scores across the four target conditions: 30, 100, 300, and 1000 targets.

## 3. Results

### Enrichments and number of targets

Each algorithm assigns an interaction score to each pair of miRNA and predicted targets. These scores help select genes that are most relevant based on predictions. In our previous work^14^, we demonstrated that selecting a few targets with the highest scores typically results in a strong enrichment score. However, including more targets may lead to less extreme enrichment scores, though this increases statistical significance due to greater statistical power. This is illustrated in Figure 1, which shows enrichment for miR-17-5p in 293T cells based on various algorithms and databases of experimentally validated targets with the top 30, 100, 300, and 1000 targets for each (see methods). To ensure a valid comparison of algorithm performance, we compared target lists of equivalent lengths for the benchmark analyses.

**Figure 1.**
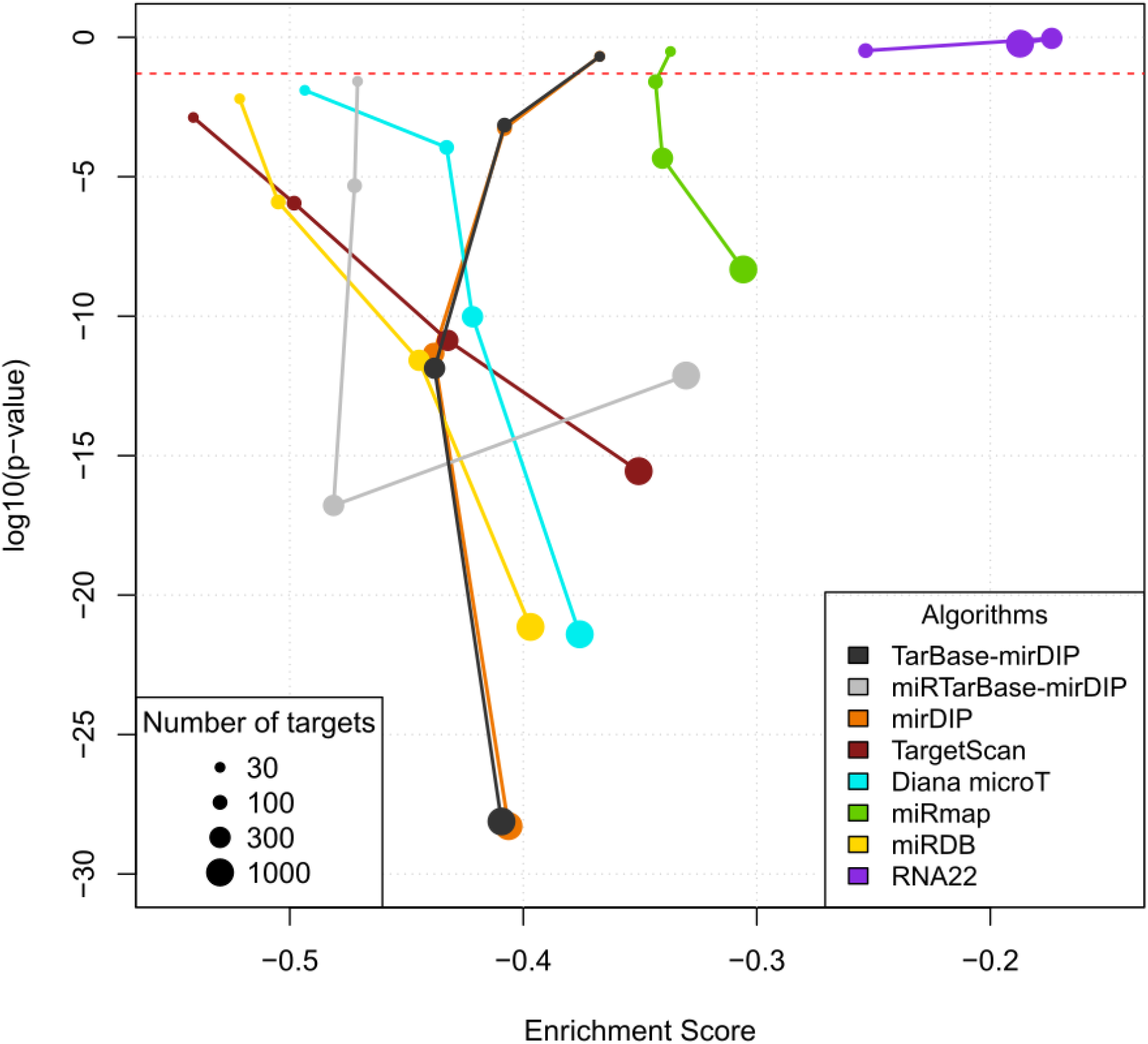
Enrichment from GSEA for miR-17-5p in 293T cells obtained with different algorithms or databases of miRNA targets. P-value is expressed in log10 scale. The analysis is performed for 30,100, 300, and 1000 top targets. For miRTarBase and TarBase the score of mirDIP was used to select the top targets. The horizontal red line stands for y =log10(0.05).

### Performances of algorithms and databases

We evaluated algorithms and databases on their ability to predict targets that are significantly anti-correlated in expression with the miRNA of interest. Our analyses were conducted on the top 30, 100, 300, and 1000 targets and we only considered miRNA with a significant negative enrichment.

Indeed, some miRNAs exhibit a significant enrichment of correlation with their predicted targets toward positive correlations. However, these unexpected cases are in the majority concerning lowly expressed or poorly variable miRNAs, which are not expected to have a strong impact on their targets. In contrast, negative enrichment was observed for highly expressed or highly variable miRNAs (Supplementary Figure 1).

We calculated the performance of the algorithms separately for four human cell lines (Figure 2) and for 22 cell types derived from mouse lung cells (Figure 3).

**Figure 2.**
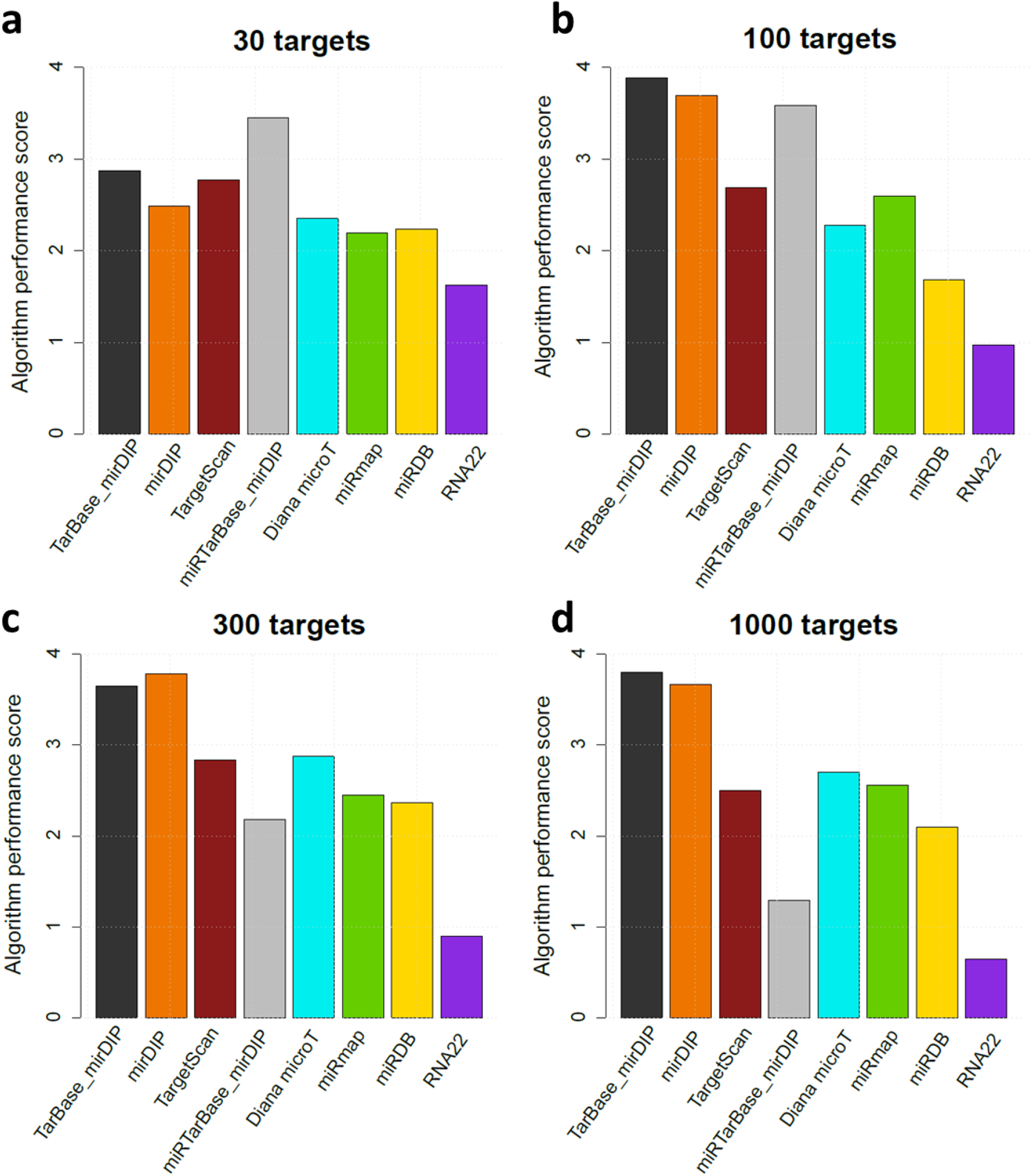
Algorithm and database performance scores for miRNA in 4 human cell lines. Scores correspond to the sum of the ratio of the number of miRNAs identified with significant negative enrichment for analysis performed for **(a)** 30, **(b)** 100, **(c)** 300, and **(d)** 1000 top targets. For miRTarBase and TarBase the score of mirDIP was used to select the top targets.

**Figure 3.**
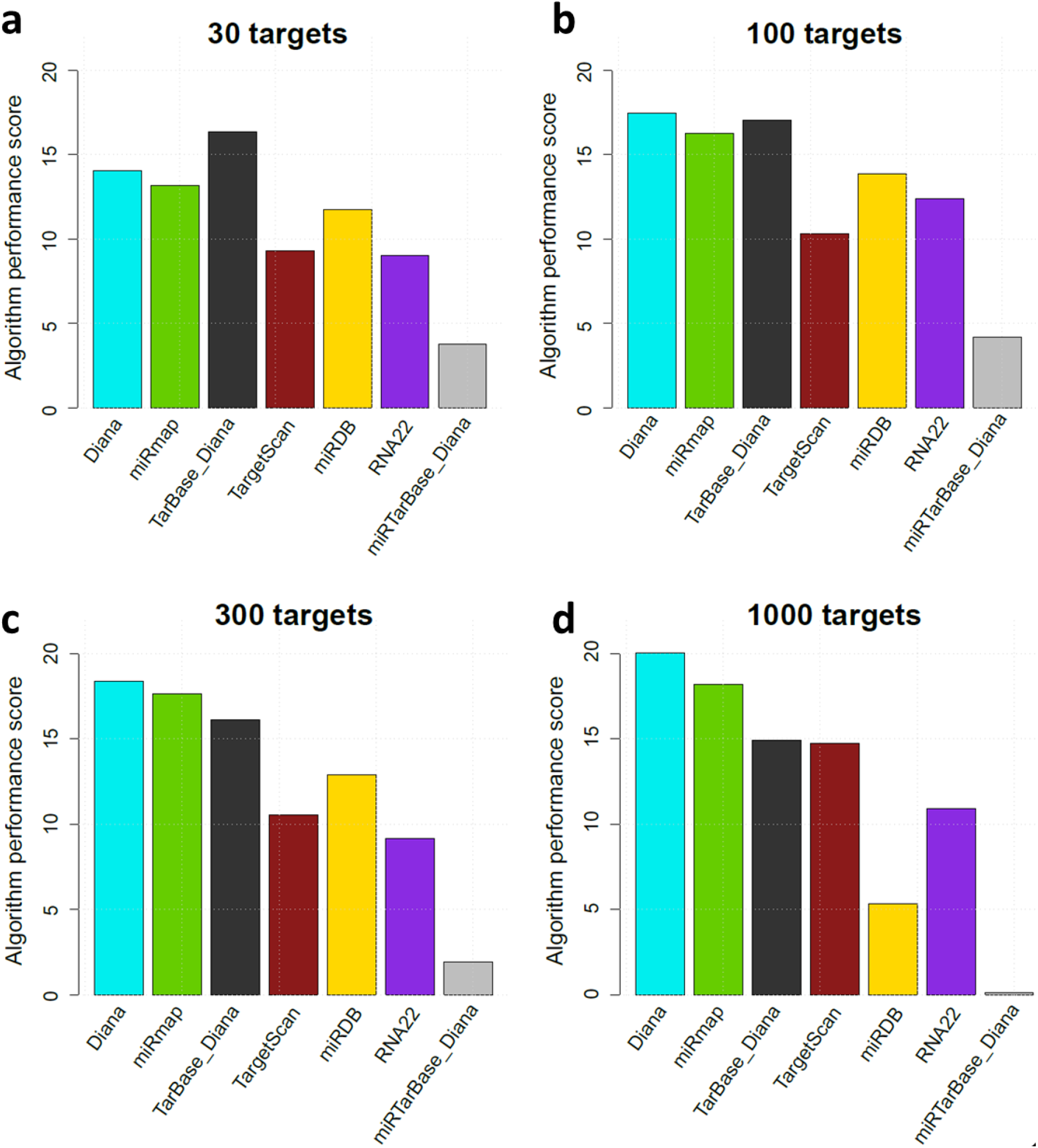
Algorithm and database performance scores for miRNA in 4 human cell lines. Scores correspond to the sum of the ratio of the number of miRNAs identified with significant negative enrichment for analysis performed for **(a)** 30, **(b)** 100, **(c)** 300, and **(d)** 1000 top targets. For miRTarBase and TarBase the score of Diana-microT was used to select the top targets.

In the analysis of human cell lines, TarBase combined with mirDIP scores and mirDIP alone demonstrated the highest overall performances. However, when focusing exclusively on 30 targets, miRTarBase, in conjunction with mirDIP scores, yielded the best results, followed by TarBase and TargetScan. Moreover, TargetScan, Diana microT, miRmap, and miRDB also showed generally strong performances, whereas RNA22 received lower scores across all conditions. It’s important to note that miRDB and miRTarBase are at a disadvantage when evaluating the top 1,000 targets due to their limited number of predicted targets; only 65% of miRNAs were tested for miRDB, and just 18% for miRTarBase.

Since mirDIP only predicts human miRNA targets, this algorithm was not included in the benchmark based on mouse cells. For mouse miRNA, Diana microT demonstrated the highest overall performance closely followed by miRmap and TarBase associated with Diana microT score. Notably, TarBase even outperformed Diana-microT for the top 30 targets. RNA22 showed overall performances similar to that of TargetScan and miRDB. The very low number of targets in miRTarBase – for which at most only 42% of miRNAs were tested for the top 30 targets – accounts for the poor performance observed. Similarly, TarBase and miRDB faced limitations by being tested on only 60% and 33 % of miRNAs for the top 1000 targets respectively.

### miRTarBase performance

In miRTarBase, experimentally validated targets do not have a numerical score but are categorized into two groups: those validated by strong evidence (such as Western-Blot, qPCR, and reporter gene assays) and those validated by weak evidence (including Microarray, NGS, pSILAC, CLIP-Seq,…). A target validated by at least one strong piece of evidence is considered a strong target.

We calculated enrichment obtained with only strong targets or with all targets for each miRNA. These results were compared to enrichments derived from using a similar number of targets (±10%) from other algorithms (Figure 4).

**Figure 4.**
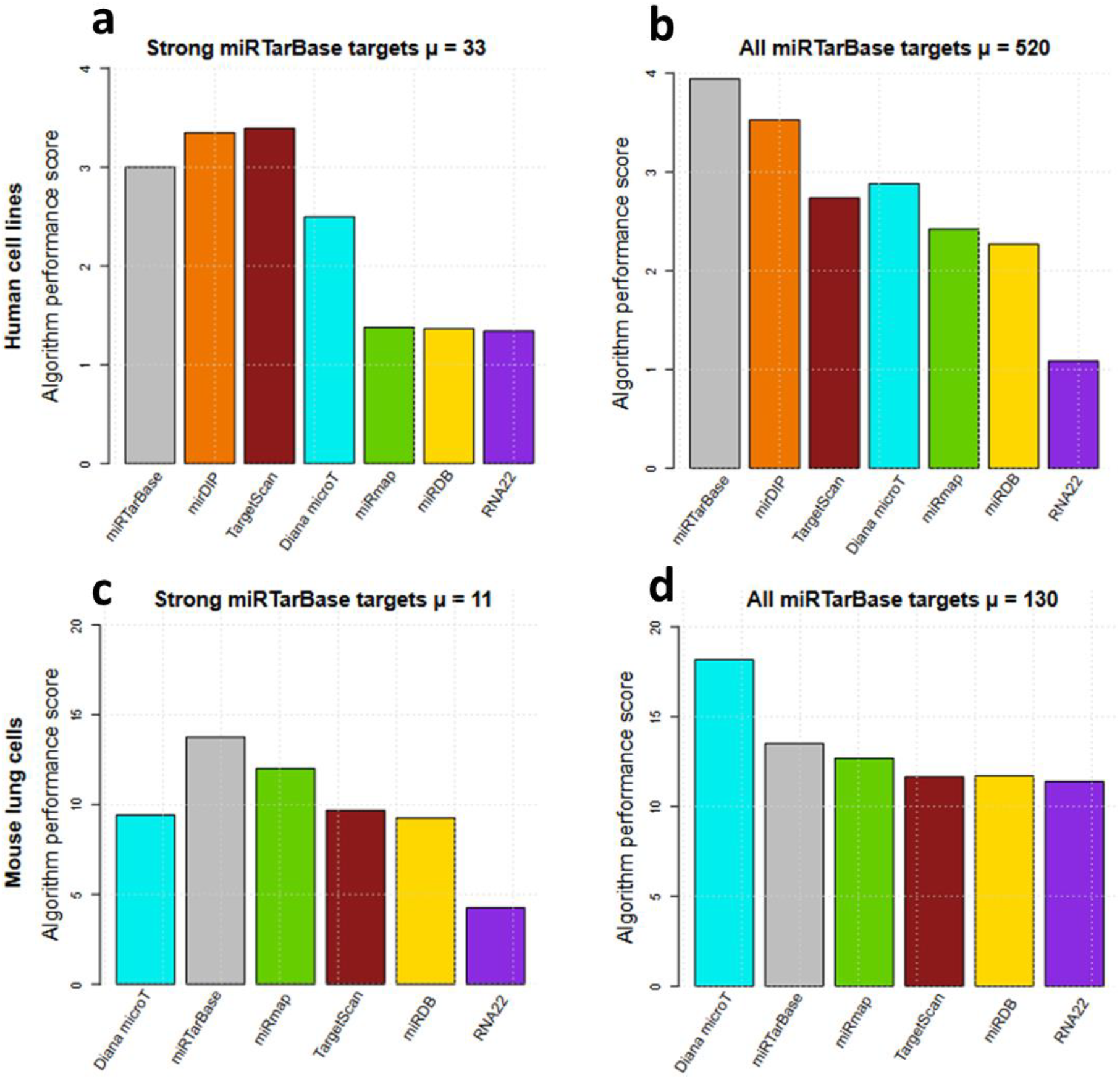
miRTarBase performance scores compare with other algorithms. Scores correspond to the sum of the ratio of the number of miRNAs identified with significant negative enrichment for analysis performed for **(a**,**b)** the 4 human cell lines and **(c**,**d)** the 22 mouse cell types. The number of targets used for each algorithm corresponds to the number of **(a**,**c)** miRTarBase strong targets, and **(b**,**d)** all targets in miRTarBase. µ corresponds to the mean number of targets used.

On average, human miRNAs have three times more strong targets and four times more overall experimentally validated targets than mouse miRNAs. In all comparisons, miRTarBase demonstrated strong performance and was the best source for all validated targets in human cells, as well as for strong targets in mouse cells. However, TargetScan and mirDIP outperformed miRTarBase for strong targets in human cells, and Diana microT performed better for an average of 130 targets in mouse cells.

### miRWalk performance

miRWalk score presents three modes, which limit flexibility regarding the number of selected targets. In order to compare miRWalk with other algorithms, a specific target count was chosen for each miRNA, aiming to get as close to 1,000 targets as possible. This same target range, adjusted by ±10%, was then applied to other algorithms if feasible. The number of targets included ranges from 95 to 6,796 for human miRNAs and from 311 to 8,367 for mouse miRNAs (Figure 5).

**Figure 5.**
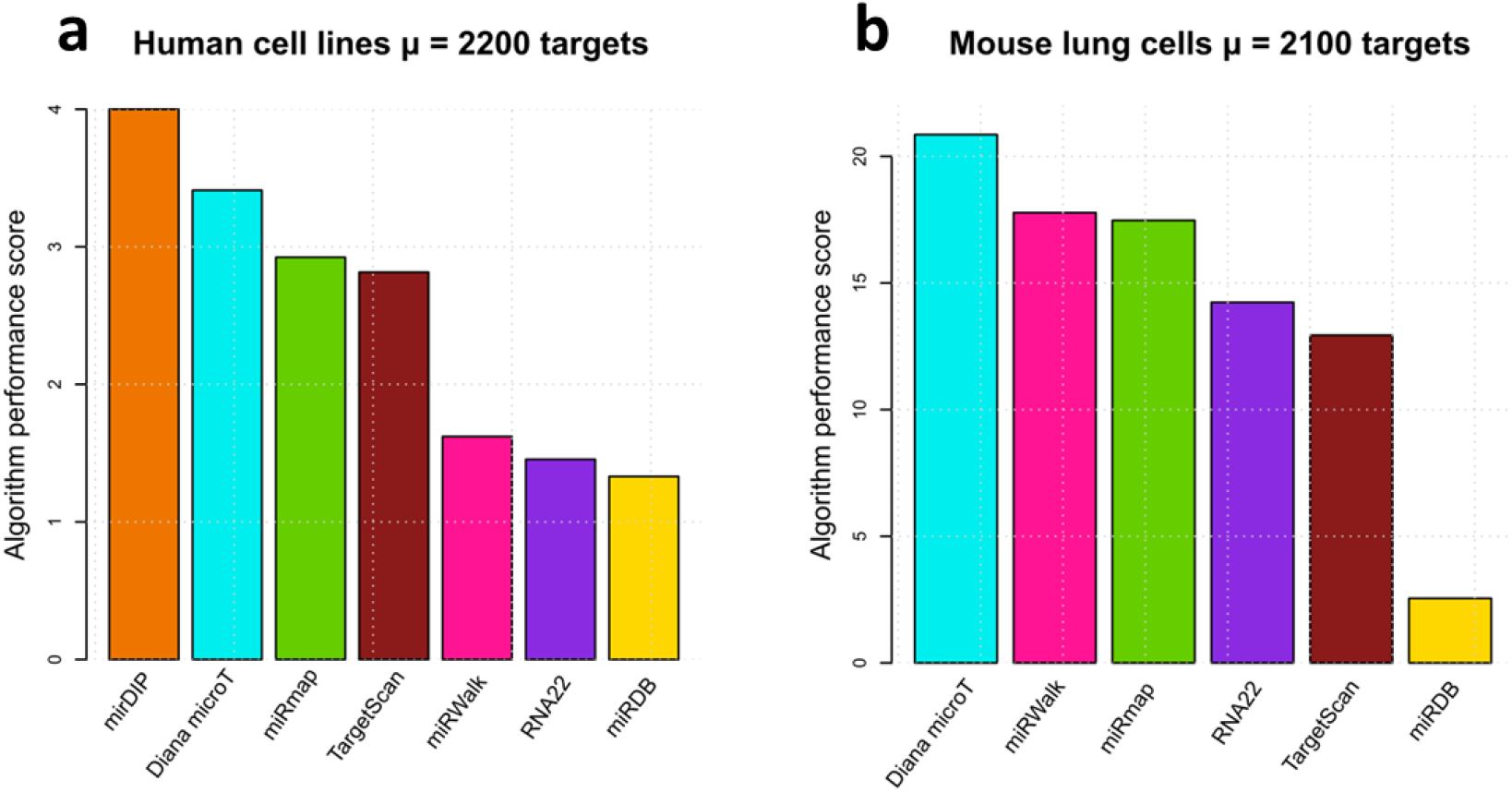
miRWalk performance scores compare with other algorithms. Scores correspond to the sum of the ratio of the number of miRNAs identified with significant negative enrichment for analysis performed for **(a)** the 4 human cell lines and **(b)** the 22 mouse cell types. The number of targets used corresponds to the number of targets selected in miRWalk to be the closest to 1000.

In both species, TargetScan and miRDB face disadvantages due to their low number of predicted targets. Consequently, only 76% of miRNAs were tested in human cells with TargetScan and only 36% with miRDB. In mouse cells, the respective percentages are 66% for TargetScan and 18% for miRDB.

In human cells, the performance of miRWalk is comparable to that of RNA22 and miRDB, although miRDB was tested on only one-third of the miRNAs. In mouse cells, miRWalk ranks just below Diana microT. However, it is important to note that miRWalk’s p-values tend to be less significant compared to those of miRmap (not shown). Due to the differences in number of predicted targets, miRWalk and miRDB are not easily comparable.

## 4. Discussion

We take advantage of exceptionally valuable datasets to study the relationship between miRNAs and their targets. We efficiently benchmark seven well-known algorithms for miRNA-target predictions, along with two databases of experimentally validated targets. This work allows us to provide clear recommendations on how to use these algorithms effectively.

To avoid bias related to enrichment variations from the number of targets used, our analyses were performed using target lists of comparable sizes.

For human cell lines, our benchmark shows high performances for mirDIP which corresponds to the consensus result of 24 different algorithms for human miRNAs. By narrowing down the miRDIP targets to those validated in TarBase, we observed a slight improvement. TargetScan, Diana microT, miRTarBase, miRmap, and miRDB, also demonstrate excellent results. However, RNA22 and miRWalk yielded poorer outcomes. When focusing on only a few dozen targets, TargetScan, along with the use of experimentally validated targets, outperformed mirDIP alone.

In mouse primary cells, Diana microT leads the performance rankings, followed closely by miRmap and TarBase when associated with the Diana microT score. Interestingly, RNA22 demonstrates better results in mouse primary cells compared to human cell lines, with its performance comparable to that of TargetScan and miRDB. Due to the limited availability of experimentally validated targets for mouse miRNAs, miRTarBase, when associated with the Diana microT score, shows poor performance. However, the use of experimentally validated targets associated with Diana microT score improved performances, for the very low number of targets – a mean of 11 for strong targets for miRTarBase and a mean of 30 targets for TarBase. miRwalk delivers relatively good results with a high number of targets in mouse cells, but its scores remain inflexible when it comes to target selection.

The strong performance demonstrated by mirDIP in human cells highlights the potential benefits of creating consensus-based tools for other species. Additionally, the improved results achieved with experimentally validated targets support the effort for conducting miRNA target validation experiments in various species.

## 5. Data Availability

Data analyzed during this study are included in the Li *et al*.^12^ article. The data of human cell lines are available in the GEO database under accession code GSE226714 [https://www.ncbi.xyz/geo/query/acc.cgi?acc=GSE226714]. The data of mouse lung cells are available on the website: https://biocaitao.github.io/PSCSRII/.

## 7. Acknowledgments

LV received funds from Ecole de l’INSERM-Pfizer Innovation (EIPI). LF received funds from the call MIC 2022 (Interdisciplinary approaches to oncogenic processes and therapeutic perspectives: Contributions to oncology of mathematics and computer science), with financial support from ITMO Cancer of Aviesan within the framework of the 2021-2030 Cancer Control Strategy, on funds administered by Inserm. The authors thank Roxane Van Olden Barneveld for her work related to the subject.

## 8. Contributions

LG designed the work, LV conducted the analysis with the help of LG, LF, and NC, LV wrote the manuscript, and all authors reviewed and agreed with the conclusions.

## 9. Competing Interests

The authors declare no competing interests.

## 10. Supplementary Figures

## Notes

### Competing Interest Statement

The authors have declared no competing interest.

## References

1. Bartel, D.P. (2018). Metazoan MicroRNAs. Cell 173, 20–51. 10.1016/j.cell.2018.03.006.

2. Tastsoglou, S., Alexiou, A., Karagkouni, D., Skoufos, G., Zacharopoulou, E., and Hatzigeorgiou, A.G. (2023). DIANA-microT 2023: including predicted targets of virally encoded miRNAs. Nucleic Acids Research 51, W148–W153. 10.1093/nar/gkad283.

3. Chen, Y., and Wang, X. (2020). miRDB: an online database for prediction of functional microRNA targets. Nucleic Acids Research 48, D127–D131. 10.1093/nar/gkz757.

4. Tokar, T., Pastrello, C., Rossos, A.E.M., Abovsky, M., Hauschild, A.-C., Tsay, M., Lu, R., and Jurisica, I. (2018). mirDIP 4.1—integrative database of human microRNA target predictions. Nucleic Acids Research 46, D360–D370. 10.1093/nar/gkx1144.

5. Vejnar, C.E., and Zdobnov, E.M. (2012). miRmap: Comprehensive prediction of microRNA target repression strength. Nucleic Acids Research 40, 11673–11683. 10.1093/nar/gks901.

6. Sticht, C., De La Torre, C., Parveen, A., and Gretz, N. (2018). miRWalk: An online resource for prediction of microRNA binding sites. PLOS ONE 13, e0206239. 10.1371/journal.pone.0206239.

7. Agarwal, V., Bell, G.W., Nam, J.-W., and Bartel, D.P. (2015). Predicting effective microRNA target sites in mammalian mRNAs. eLife 4. 10.7554/eLife.05005.

8. Miranda, K.C., Huynh, T., Tay, Y., Ang, Y.-S., Tam, W.-L., Thomson, A.M., Lim, B., and Rigoutsos, I. (2006). A Pattern-Based Method for the Identification of MicroRNA Binding Sites and Their Corresponding Heteroduplexes. Cell 126, 1203–1217. 10.1016/j.cell.2006.07.031.

9. Cui, S., Yu, S., Huang, H.-Y., Lin, Y.-C.-D., Huang, Y., Zhang, B., Xiao, J., Zuo, H., Wang, J., Li, Z., et al. (2025). miRTarBase 2025: updates to the collection of experimentally validated microRNA–target interactions. Nucleic Acids Research 53, D147–D156. 10.1093/nar/gkae1072.

10. Skoufos, G., Kakoulidis, P., Tastsoglou, S., Zacharopoulou, E., Kotsira, V., Miliotis, M., Mavromati, G., Grigoriadis, D., Zioga, M., Velli, A., et al. (2024). TarBase-v9.0 extends experimentally supported miRNA–gene interactions to cell-types and virally encoded miRNAs. Nucleic Acids Research 52, D304–D310. 10.1093/nar/gkad1071.

11. Subramanian, A., Tamayo, P., Mootha, V.K., Mukherjee, S., Ebert, B.L., Gillette, M.A., Paulovich, A., Pomeroy, S.L., Golub, T.R., Lander, E.S., et al. (2005). Gene set enrichment analysis: A knowledge-based approach for interpreting genome-wide expression profiles. Proceedings of the National Academy of Sciences 102, 15545–15550. 10.1073/pnas.0506580102.

12. Li, J., Tian, J., and Cai, T. (2025). Integrated analysis of miRNAs and mRNAs in thousands of single cells. Scientific Reports 15. 10.1038/s41598-025-85612-z.

13. Yu, G. (2024). Thirteen years of clusterProfiler. The Innovation 5, 100722. 10.1016/j.xinn.2024.100722.

14. Velut, L. (2024). Single-cell microRNA-mRNA co-sequencing techniques convey large potential for understanding microRNA regulations but require careful and systemic approaches, IRIG-BCI-IMAC/Dual_miR_mRNA_SCseq. Version Version1 (Zenodo). 10.5281/ZENODO.14516469 10.5281/ZENODO.14516469.

