## Supplementary Figure 1 for "Benchmarking microRNA Target Prediction Algorithms Using Single-Cell Co-Sequencing Data"

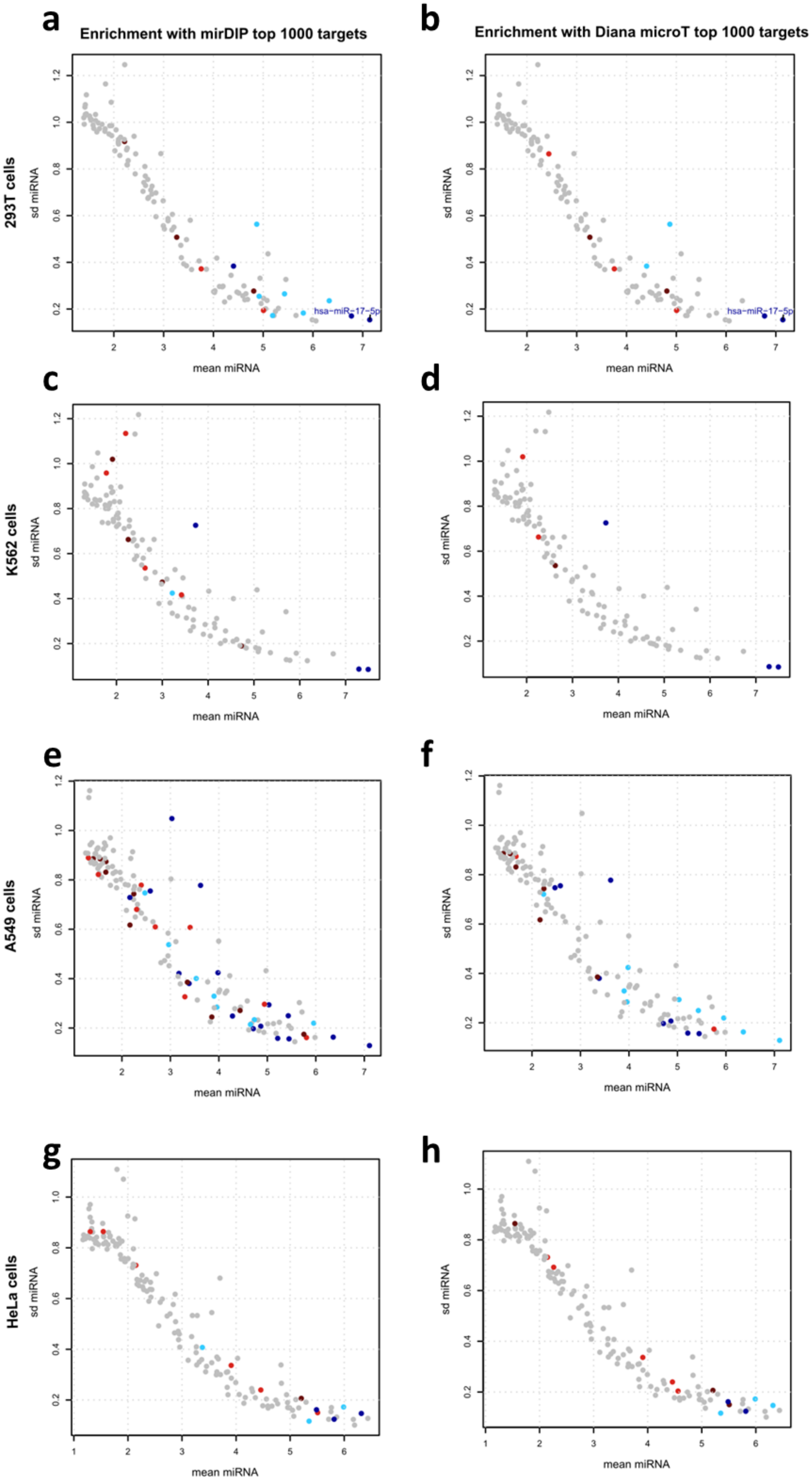
a
Enrichment with mirDIP top 1000 targets
b
Enrichment with Diana microT top 1000 targets
293T cells
sd miRNA
mean miRNA
hsa-miR-17-5p
c
d
K562 cells
e
f
A549 cells
g
h
HeLa cells

**Supplementary Figure 1. Standard deviation by means of miRNA expression colored by GSEA results for human cell lines.** Panels correspond to **(a,b)** 293T cells, **(c,d)** K562 cells, **(e,f)** A549 cells, and **(g,h)** HeLa cells. GSEA was computed with the top 1,000 targets predicted by **(a,c,e,g)** mirDIP and **(b,d,f,h)** Diana microT. Each point represents a miRNA. miRNAs with no significant enrichment are represented in grey. miRNAs with a significant enrichment toward negative (resp. positive) correlation are represented in blue (resp. red). A darker color corresponds to a p-value lower than 0.005.
